# Coevolutive, Evolutive and Stochastic Information in Protein-Protein Interactions

**DOI:** 10.1101/683128

**Authors:** Miguel Andrade, Camila Pontes, Werner Treptow

**Author notes:** Corresponding Author: Werner Treptow, Laboratório de Biologia Teórica e Computacional (LBTC), Universidade de Brasília DF, Brazil. **Author contributions** WT designed research; MA and CP performed research; MA, CP and WT analyzed data; WT wrote original draft; WT, MA and CP reviewed and edited. MA and CP contributed equally to this work.

## Abstract

Here, we investigate the contributions of coevolutive, evolutive and stochastic information in determining protein-protein interactions (PPIs) based on primary sequences of two interacting protein families *A* and *B*. Specifically, under the assumption that coevolutive information is imprinted on the interacting amino acids of two proteins in contrast to other (evolutive and stochastic) sources spread over their sequences, we dissect those contributions in terms of compensatory mutations at physically-coupled and uncoupled amino acids of *A* and *B*. We find that physically-coupled amino-acids at short range distances store the largest per-contact mutual information content, with a significant fraction of that content resulting from coevolutive sources alone. The information stored in coupled amino acids is shown further to discriminate multi-sequence alignments (MSAs) with the largest expectation fraction of PPI matches – a conclusion that holds against various definitions of intermolecular contacts and binding modes. When compared to the informational content resulting from evolution at long-range interactions, the mutual information in physically-coupled amino-acids is the strongest signal to distinguish PPIs derived from cospeciation and likely, the unique indication in case of molecular coevolution in independent genomes as the evolutive information must vanish for uncorrelated proteins.

**SIGNIFICANCE:** The problem of predicting protein-protein interactions (PPIs) based on multi-sequence alignments (MSAs) appears not completely resolved to date. In previous studies, one or more sources of information were taken into account not clarifying the isolated contributions of coevolutive, evolutive and stochastic information in resolving the problem. By benefiting from data sets made available in the sequence- and structure-rich era, we revisit the field to show that physically-coupled amino-acids of proteins store the largest (per contact) information content to discriminate MSAs with the largest expectation fraction of PPI matches – a result that should guide new developments in the field, aiming at characterizing protein interactions in general.

## INTRODUCTION

While being selected to be thermodynamically stable and kinetically accessible in a particular fold (1, 2), interacting proteins *A* and *B* coevolve to maintain their bound free-energy stability against a vast repertoire of non-specific partners and interaction modes. Protein coevolution, in the form of a time-dependent molecular process, then translates itself into a series of primary-sequence variants of *A* and *B* encoding coordinated compensatory mutations (3) and, therefore, specific protein-protein interactions (PPIs) derived from this stability-driven process (4). As a ubiquitous process in molecular biology, coevolution thus apply to protein interologs, either paralogous or orthologous, under cospeciation or in independent genomes.

Thanks to extensive investigations in the past following ingenious approaches based on the correlation of phylogenetic trees (5–7) and profiles (8), gene colocalization (9) and fusions (10), maximum coevolutionary interdependencies (11) and correlated mutations (12, 13), the problem of predicting PPIs based on multi-sequence alignments (MSAs) appears to date resolvable, at least for small sets of paralogous sequences - recent improvements (14–18) resulting from PPI prediction allied to modern coevolutionary approaches (19–23) to identify interacting amino acids across protein interfaces. In these previous studies, however, the information was taken into account as a whole, and it was not clarified, as discussed in recent reviews (4, 24), the isolated contributions of coevolutive, evolutive and stochastic information in resolving the problem.

Here, by benefiting from much larger datasets made available in the sequence- and structure-rich era, we revisit the field by quantifying the amount of information that protein *A* stores about protein *B* stemming from each of these sources and, more importantly, their effective contributions in determining PPIs based on MSAs (Scheme 1). Specifically, under the assumption that the coevolutive information is imprinted on the interacting amino acids of protein interologs in contrast to other (evolutive and stochastic) sources spread over their sequences, we want that information to be dissected in terms of compensatory mutations at physically-coupled and uncoupled amino acids of *A* and *B*. Anticipating our findings, we show that physically-coupled amino-acids store the largest per-contact mutual information (MI) content to discriminate concatenated MSAs with the largest expectation fraction of PPI matches – a conclusion that holds against various definitions of intermolecular protein contacts and binding modes. A significant fraction of that information results from coevolutive sources alone. It is important to acknowledge that, even though our findings are derived for a set of interologs under cospeciation, they can be generalized to cases of non-cospeciating interologs given that the underlying thermodynamical principles must be the same for all cases.

**Scheme 1.**
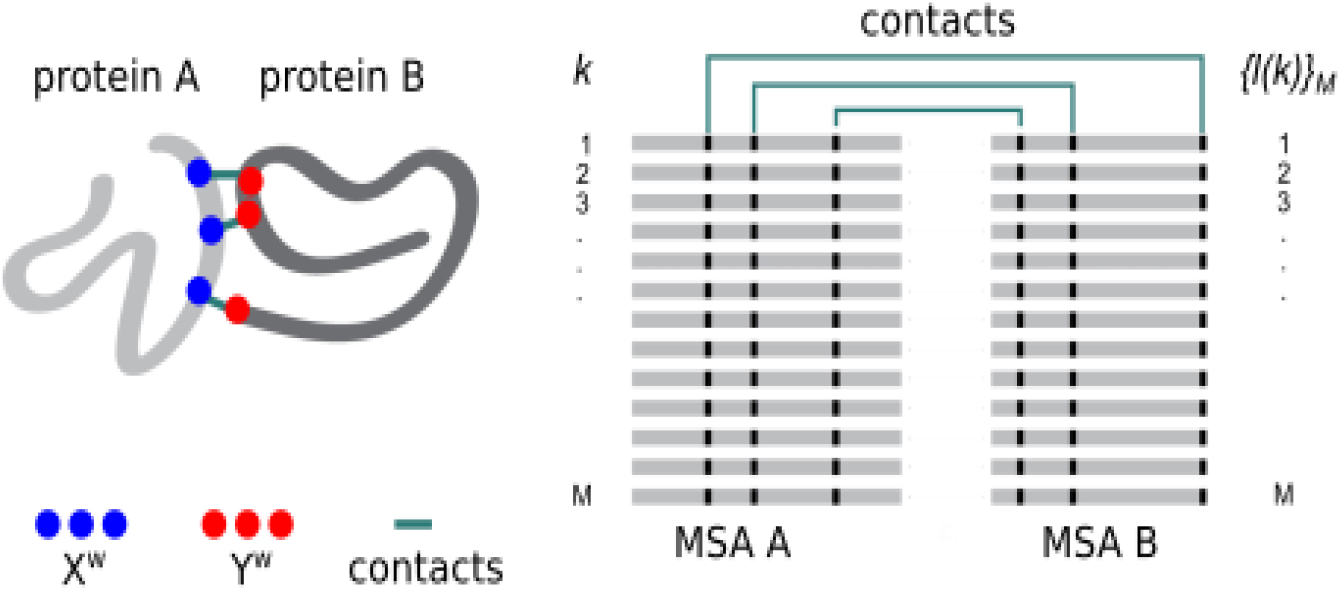
Structural contacts mapped into M-long multi-sequence alignment (MSA) of protein interologs A and B. A set of pairwise protein-protein interactions is defined by associating each sequence *l* in MSA *B* to a sequence *k* in MSA *A* in one unique arrangement, {*l* (*k*)∣*z*}_*M*_, determined by the coevolution process *z* to which these protein families were subjected.

## THEORY AND METHODS

### Decomposition of Mutual Information

In detail, consider two proteins *A* and *B* that interact via formation of *i*=1,…, *N* amino-acid contacts at the molecular level. Proteins *A* and *B* are assumed to coevolve throughout *M !* distinct processes *z* described by the stochastic variable *Z* with an uniform probability mass function ρ(*z*), ∀ *z* ∈{1,…, *M !*}. Given any specific process *z*, their interacting amino-acid sequences are respectively described by two *N*-length blocks of discrete stochastic variables *X*^*N*^≡(*X*_1_,…, *X*_*N*_) and *Y*^*N*^≡(*Y*_1_,…, *Y*_*N*_) with probability mass functions {ρ(*x*^*N*^),ρ(*y*^*N*^),ρ(*x*^*N*^, *y*^*N*^∣*z*)} such that,

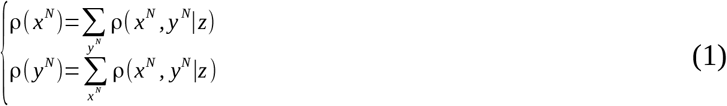

and

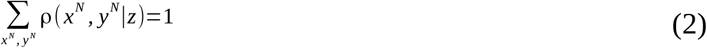

for every joint sequence {*x*^*N*^, *y*^*N*^}_|χ|^*2N*^_ defined in the alphabet χ of size |χ|. Under these considerations, the amount of information that protein *A* stores about protein *B* is given by the mutual information *I*(*X*^*N*^; *Y*^*N*^∣*z*) between *X*^*N*^ and *Y*^*N*^ conditional to process *z* (25). As made explicit in eq. [1], we are particularly interested in quantifying *I* (*X*^*N*^; *Y*^*N*^∣*z*) for the situation in which marginals of the *N*-block variables {ρ(*x*^*N*^),ρ(*y*^*N*^)} are assumed to be independent of process *z* meaning that, for a fixed sequence composition of proteins *A* and *B* only their joint distribution depends on the process. Furthermore, by assuming *N*-independent contacts, we want that information to be quantified for the least-constrained model ρ^*^(*x*^*N*^, *y*^*N*^∣*z*) that maximizes the conditional joint entropy between *A* and *B* - that condition ensures the mutual information to be written exactly, in terms of the individual contributions of contacts *i*.

For the least-constrained distribution { ρ^*^(*x*^*N*^, *y*^*N*^∣*z*)}, the conditional mutual information

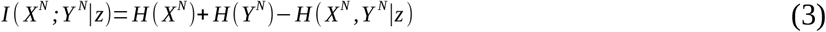

writes in terms of the Shannon’s information entropies

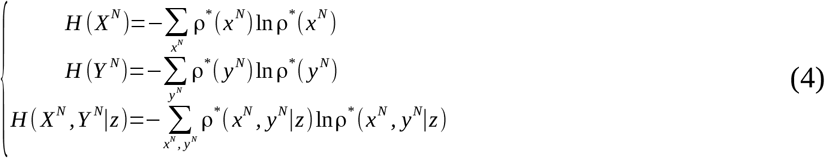

associated with the conditional joint distribution {ρ^*^(*x*^*N*^, *y*^*N*^∣*z*)} and the derived marginals {ρ^*^(*x*^*N*^), ρ^*^(*y*^*N*^)} of the *N*-block variables. From its entropy-maximization property, the critical distribution ρ^*^(*x*^*N*^, *y*^*N*^∣*z*) factorizes into the conditional two-site marginal of every contact *i*

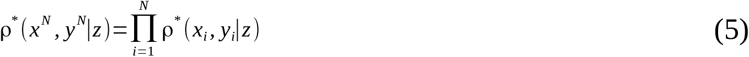

then allowing eq.[4] to be written extensively, in terms of the individual entropic contributions

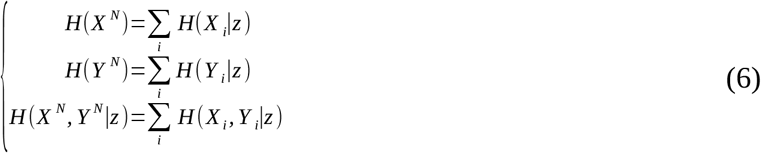

such that,

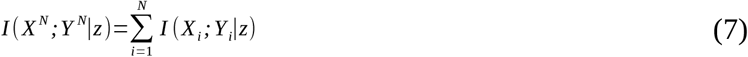

(cf. *SI* for details). In eq. [7], the conditional mutual information achieves its lower bound of zero if *X*^*N*^ and *Y*^*N*^ are conditionally independent given *z ie.*, ρ^*^(*x*^*N*^, *y*^*N*^∣*z*)=ρ^*^(*x*^*N*^)×ρ^*^(*y*^*N*^). For the case of perfectly correlated variables ρ^*^(*x*^*N*^, *y*^*N*^∣*z*)=ρ^*^(*x*^*N*^)=ρ^*^(*y*^*N*^), the conditional mutual information is bound to a maximum which cannot exceed the entropy of either block variables *H* (*X*^*N*^) and *H* (*Y*^*N*^).

Given a known set of protein amino-acid contacts and their underlying primary sequence distributions defining the stochastic variables *X*^*N*^ and *Y*^*N*^, eq. [7] thus establishes the formal dependence of their mutual information with any given process *z*. Because “contacts” can be defined for a variety of cutoff distances *r*_*c*_, eq. [7] is particularly useful to dissect mutual information in terms of physically-coupled and uncoupled protein amino acids. In the following, we explore eq. [7] in that purpose by obtaining the two-site probabilities in eq. [5]

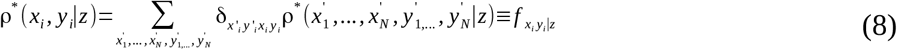

from the observed frequencies 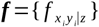 in the multiple-sequence alignment

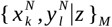

where the *N*-length amino-acid block *l* of protein *B* is joint to block *k* of protein *A* in one unique arrangement { *l* (*k*)∣*z*}_*M*_ for 1≤*k* ≤ *M* (*cf.* Scheme 1 and Computational Methods).

### Computational Methods

Table-S1 details the interacting protein systems considered in the study. For each system under investigation, amino-acid contacts defining the discrete stochastic variables *X*^*N*^ and *Y*^*N*^ including physically coupled amino acids at short-range cut-off distances (*r*_*c*_≤8.0 Å) and physically uncoupled amino-acids at long-range cut-off distances (*r*_*c*_> 8.0 Å) were identified from the x-ray crystal structure of the bound state of proteins *A* and *B*. The reference (native) multi-sequence alignment 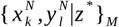 of the joint amino-acid blocks associated to *X*^*N*^ and *Y*^*N*^ was reconstructed from annotated primary-sequence alignments published by Baker and coworkers (22), containing *M* paired sequences with known PPI interactions and defined in the alphabet of 20 amino acids plus the gap symbol (_|χ|_=21). Scrambled MSA models were generated by randomizing the pattern {*l* (*k*)∣*z* ^*^}_*M*_ in which block *l* is joint to block *k* in the reference alignment.

For any given MSA model, two-site probabilities 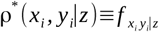 were defined from the observable frequencies 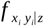 regularized by a pseudocount effective fraction λ^*^ in case of insufficient data availability as devised by Morcos and coauthors (19). More specifically, two-site frequencies were calculated according to

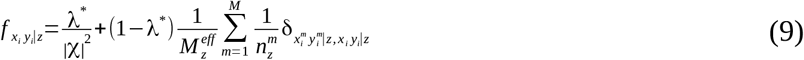

where, 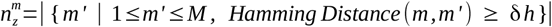 is the number of similar sequences *m*′ within a certain Hamming distance δ*h* of sequence *m* and 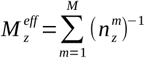 is the effective number of distinguishable primary sequences at that distance threshold - the Kronecker delta 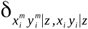 ensures counting of (*x*_*i*_, *y*_*i*_) occurrences only. In eq. [9], two-site frequencies converge to raw occurrences in the sequence alignment for λ^*^=0 or approach the uniform distribution 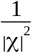 for λ^*^=1; eq. [9] is identical to the equation devised by Morcos and coauthors (19) by rewriting 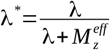. Here, two-site probabilities 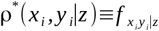 were computed from eq. [9] after unbiasing the reference MSA by weighting down primary sequences with amino-acid identity equal to 100%. An effective number of primary sequences 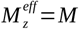 (*cf.* Table-S1) was retained for analysis and a pseudocount fraction of λ^*^=0.001 was used to regularize data without largely impacting observable frequencies. Single-site probabilities { ρ(*x*^*N*^),ρ(*y*^*N*^)} were derived from ρ^*^(*x*_*i*_, *y*_*i*_∣*z*) by marginalization via eq. [1].

The conditional mutual information in eq. [7] was computed from single- and joint-entropies according to eq. [3]. Given the fact that the maximum value of *I* (*X*_*i*_; *Y*_*i*_∣*z*) is bound to the conditional joint entropy, eq. [7] was computed in practice as a per-contact entropy-weighted conditional mutual information (26), *H*(*X*_*i*_,*Y*_*i*_∣*z*)^−1^ *I* (*X*_*i*_; *Y*_*i*_∣*z*), to avoid that contributions of contacts between highly variable sites are overestimated. Because *H*(*X*_*i*_,*Y*_*i*_∣*z*) and *I* (*X*_*i*_; *Y*_*i*_∣*z*) have units of *nats*, eq. [7] is dimensionless in the present form.

## RESULTS AND DISCUSSION

Details of all protein systems under investigation are presented in Table-S1. Each system involves two families of protein interologs *A* and *B* with known PPIs derived from cospeciation in the same genome (27). We denote by 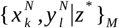 their reference concatenated MSA associated to the native process *z* ^*^. For convenience, in the following, we present and discuss results obtained for a representative system *A* and *B* – the protein complex TusBCD (chains B and C of 2DIP) which is crucial for tRNA modification in *Escherichia coli*. Similar results and conclusions hold for all other systems in Table-S1 as presented in supplementary figures S1 through S4 (*cf.* SI).

### Decomposition of Mutual Information

Fig. 1A shows the three-dimensional representation of variables *X*^*N*^ and *Y*^*N*^ embodying every possible amino-acid pairs along proteins *A* and *B* and their decomposition in terms of physically coupled amino acids at short-range cutoff distances (*r*_*c*_≤8.0 Å) and physically uncoupled amino-acids at long-range cutoff distances (*r*_*c*_>8.0 Å). In Fig. 1B, the total mutual information *I* (*X*^*N*^; *Y*^*N*^∣*z*^*^) across every possible amino-acid pairs of *A* and *B* (coupled + uncoupled) amounts to 987.88 in the reference (native) MSA (process *z* ^*^). As estimated from a generated ensemble of scrambled MSA models, expectation values for the mutual information 〈 *I*(*X*^*N*^; *Y*^*N*^∣*z*)〉_*M* − *n*_ decreases significantly as decorrelation or the number *M* − *n* of mismatched proteins in the reference MSA increases, for 0≤ *n*≤*M*. The result also holds at the level of individual protein contacts *i* as *I* (*X*_*i*_; *Y*_*i*_∣*z*^*^) for the reference alignment is systematically larger than 〈 *I*(*X*_*i*_; *Y*_*i*_∣*z*)〉_*M*− *n*_ for scrambled MSA models with *M* sequence mismatches (Fig. S1).

**Figure 1.**
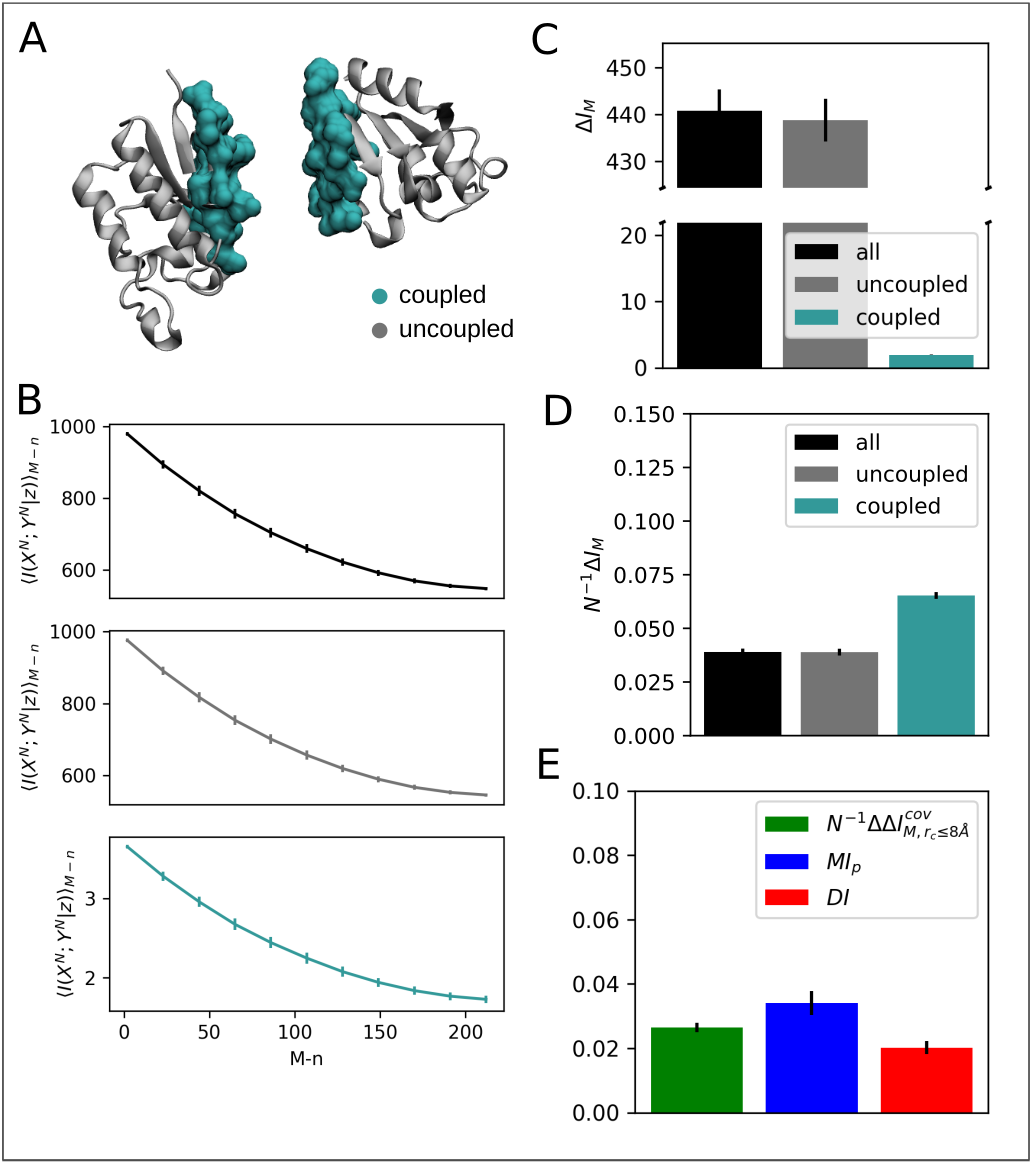
Informational analysis of protein complex TusBCD, chains *B* and *C*. (A) Three-dimensional representation of stochastic variables *X*^*N*^ and *Y*^*N*^ as defined from physically coupled amino acids at short-range cutoff distances *r*_*c*_≤8.0 Å (turquoise) and physically uncoupled amino-acids at long-range cutoff distances *r*_*c*_>8.0 Å (gray). (B) Conditional mutual information 〈*I*(*X*^*N*^;*Y*^*N*^ ∣*z*)〉_*M* − *n*_ as a function of the number *M* − *n* of randomly paired proteins in the reference (native) MSA. 〈*I*(*X*^*N*^;*Y*^*N*^ ∣*z*)〉_*M* − *n*_ are expectation values estimated from a generated ensemble of 500 MSA models. (C) Mutual information gap Δ *I*_*M*_ between reference and 100 random models featuring *M* randomly paired sequences. (D) Per-contact mutual information gap 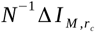. (E) Mutual information decomposition 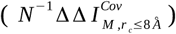 according to eq. [11] and comparison with functional mutual information 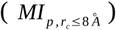 and direct information 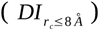. In B, C, D and E error bars correspond to standard deviations.

As a measure of correlation, it is not surprising that mutual information in the reference MSA is larger than that of scrambled alignments. Not expected, however, is the fact that correlation does not vanish at scrambled models meaning that part of the calculated mutual information results at random. Supporting that notion, low mutual information between amino acid positions of multiple sequence alignments are typically indicative of independently evolved proteins. Subtraction of that stochastic source from *I* (*X*^*N*^; *Y*^*N*^∣*z*^*^) as computed in the form of a mutual-information gap

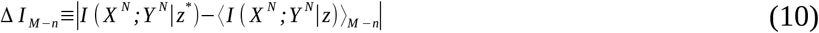

between the reference model and models with *M* mismatched sequences then reveals the isolated nonstochastic contributions to the total correlation between proteins *A* and *B*. Here, the information gap Δ *I*_*M*_ amounts to ~ 440 for every possible amino-acid pairs along proteins *A* and *B*. Fig. 1C shows the individual contributions of physically coupled and uncoupled amino acids to the total mutual information gap, 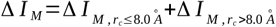. As a direct consequence of the extensive property of eq. [7], 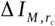 increases with *r*_*c*_ and, consequently, with the block length *N* of the corresponding stochastic variables. As such, the information imprinted at physically uncoupled amino acids accounts for most of the total mutual information gap (438.8132 ± 4.5159). When normalized by the block length or the number of contacts (Fig. 1D), 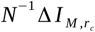, reveals a distinct dependence being larger for physically coupled amino acids than uncoupled ones (0.0653 ± 0.0015 *versus* 0.039 ± 0.0004). The information gap profile as a function of amino-acid pair distances shown in Fig. S2 makes sense of the result by showing few larger values of Δ *I*_*M*_ at short amino-acid pair distances in contrast to many smaller ones at long distances.

Under the assumption that the coevolutive information is imprinted on the interacting amino acids of interologs in contrast to other (evolutive and stochastic) sources spread over their primary sequences, the difference between short- and long-range contributions provides us with per-contact estimates for the information content resulting from coevolution alone that is,

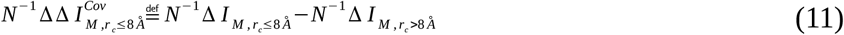

where, 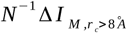 represents the per-contact mutual information resulting from evolution. As shown in Fig. 1E, 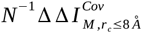 amounts to 0.0264 ± 0.0014 which compares well to independent measures of coevolutionary information *ie.*, functional mutual information 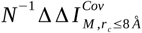(26) and direct information 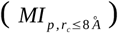 (19), 0.0340 ± 0.0037 and 0.0202 ± 0.0019. More specifically, *MI*_*p*_ is a metric 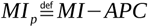 formulated by Dunn and coworkers (26) in which mutual information is subtracted from structural or functional relationships (*APC*) whereas, *DI* is based on the direct coupling analysis that removes all kinds of indirect correlations by following a global statistical approach (19). According to definition in eq. [11], we then conclude that ~ 40 % of the information content stored in physically coupled amino acids of the protein complex TusBCD results from coevolutive sources alone.

### Degeneracy and Error Analysis of Short and Long-Range Correlations

The present analysis reveals quantitative differences between short- and long-range correlations of proteins *A* and *B*. Because 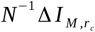 provides us with an unbiased (intensive) estimate for proper comparison of the information contents between coupled and uncoupled amino acids, in the following, we focus our attention on 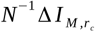 to dissect their effective contributions in determining PPIs based on sequence alignments. Accordingly, let us define the total number ω_*S*_ of native-like MSA models generated by randomization of *M* − *n* sequence pairs in the reference alignment

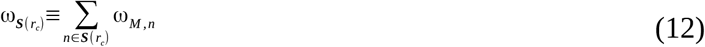

in terms of *rencontres* numbers ω_*M,n*_

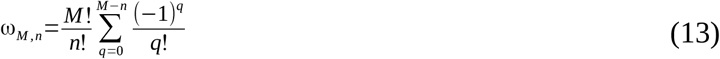

or permutations of the reference sequence set {*l* (*k*)∣*z* ^*^}_*M*_ with *n* fixed positions satisfying 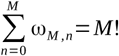 (in combinatorics language). Here, ***S***(*r*_*c*_) denotes the set of fixed positions *n*

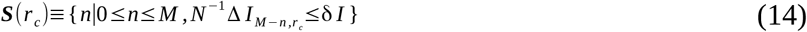

for which the normalized mutual information gap 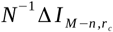, is smaller than a certain resolution δ*I* independently from the corresponding block length *N* or the number of amino-acid contacts. In simple terms, ω_*S*_ informs us on the degeneracy or the number of scrambled MSA models with a similar amount of mutual information of that in the reference multi-sequence alignment.

As shown in Table-S2, ω_*M,n*_ is an astronomically increasing function of *M* − *n*, identical for any definition of *X*^*N*^ and *Y*^*N*^ derived from the same number *M* of aligned sequences. For instance, there is 164548102752 alignments for the protein complex TusBCD with *M* − *n*=5 scrambled sequence pairs. In contrast, ω_*S*_ depends on the stochastic variables at various resolutions δ*I*(Fig. 2A). The number of native-like MSA models is substantially smaller for definitions of *X*^*N*^ and *Y*^*N*^ embodying physically-coupled amino acids in consequence of the smaller number *M* − *n* of randomly paired sequences required to perturb 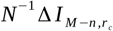, of a fixed change δ *I* such that ω_*S*_ accumulates less over MSA models satisfying 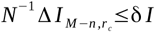(Fig. 2B).

**Figure 2.**
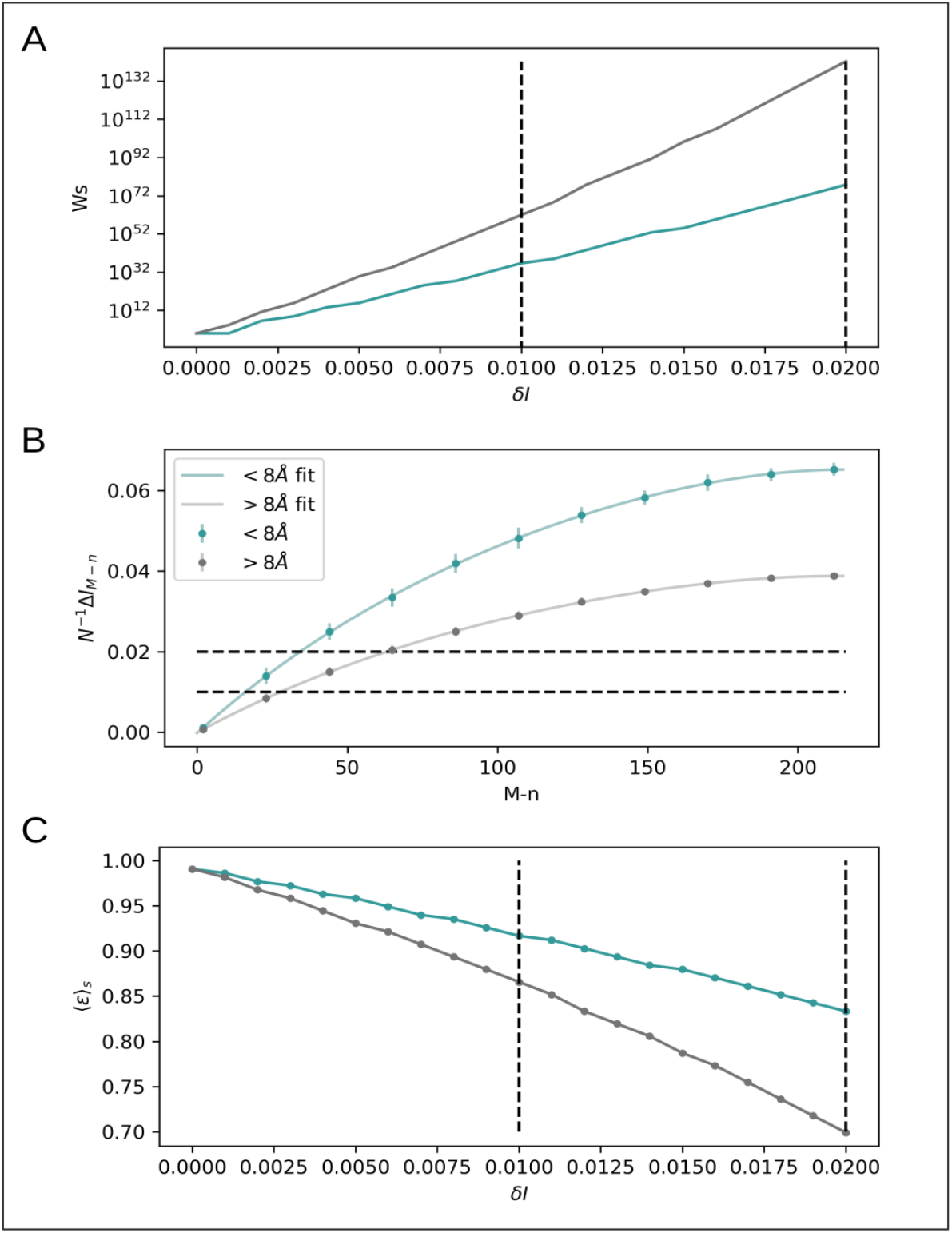
Degeneracy and error analysis for *X*^*N*^ and *Y*^*N*^ involving interacting amino acids at short-range distances *r*_*c*_≤8.0 Å (turquoise) and long-range distances *r*_*c*_>8.0 Å(gray). (A) Total number ω_*S*_ of native-like MSA models at various resolutions δ *I*. (B) Per-contact gaps of mutual information 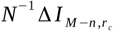, as a function of the number *M* − *n* of scrambled sequence pairs in the reference alignment. (C) Expectation values 〈ϵ〉_*S*_ (eq. [15]) for the fraction of sequence matches across native-like MSA models at various resolutions δ *I*. Dashed lines highlight differences at δ *I* values of 0.01 and 0.02.

The degeneracy of *M*-long native-like MSA models at a given resolution δ *I* depends on the cutoff distance *r*_*c*_ defining *X*^*N*^ and *Y*^*N*^ (Fig. 2A). That condition imposes distinct boundaries for the amount of PPIs amenable of resolution across definitions of the stochastic variables in terms of coupled and uncoupled amino acids. Indeed, the expectation value

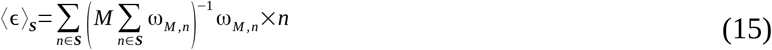

for the fraction *M*^−1^*n* of primary sequence matches among native-like MSA models mapped by *S* in eq. [14] decreases substantially with ω_*S*_ meaning that 〈ϵ〉_*S*_ is systematically larger for physically-coupled amino-acids at various resolutions δ *I* (Fig. 2C). For instance, the fraction of matches at δ *I* =0.02 is ~ 20% larger for coupled amino-acids than the same estimate for amino acids at long-range distances (0.8333 *versus* 0.6991). Linear extrapolation in Fig. 2C along increased values of δ *I* suggests even larger differences in the expectation fraction of PPI matches between short and long-range correlations of proteins *A* and *B*.

### Dependence with Contact Definition 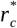 and Docking Decoys

So far, our results were determined by defining physically coupled amino acids at short-range cutoff distances 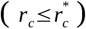 and physically uncoupled amino-acids at long-range cutoff distances 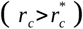, for 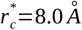. Further analysis shows however a clear dependence of the per-contact mutual information gap 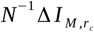, of coupled amino acids with 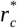 – which is not the case for uncoupled ones. As shown in Fig. 3A that distinction is due the coevolutive information stored at short-range distances which reaches a maximum at 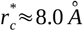 in contrast to evolutive sources uniformly spread over an entire range of 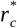 values. Particularly interesting the result strongly support the assumption that coevolutive information is imprinted preferentially on physically-coupled amino acids of interologs in contrast to other (evolutive and stochastic) sources spread over their primary sequences - a conclusion further supported by calculations of the mutual information *MI*_*p*_ subtracted from structural or functional relationships as a function of 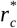.

**Figure 3.**
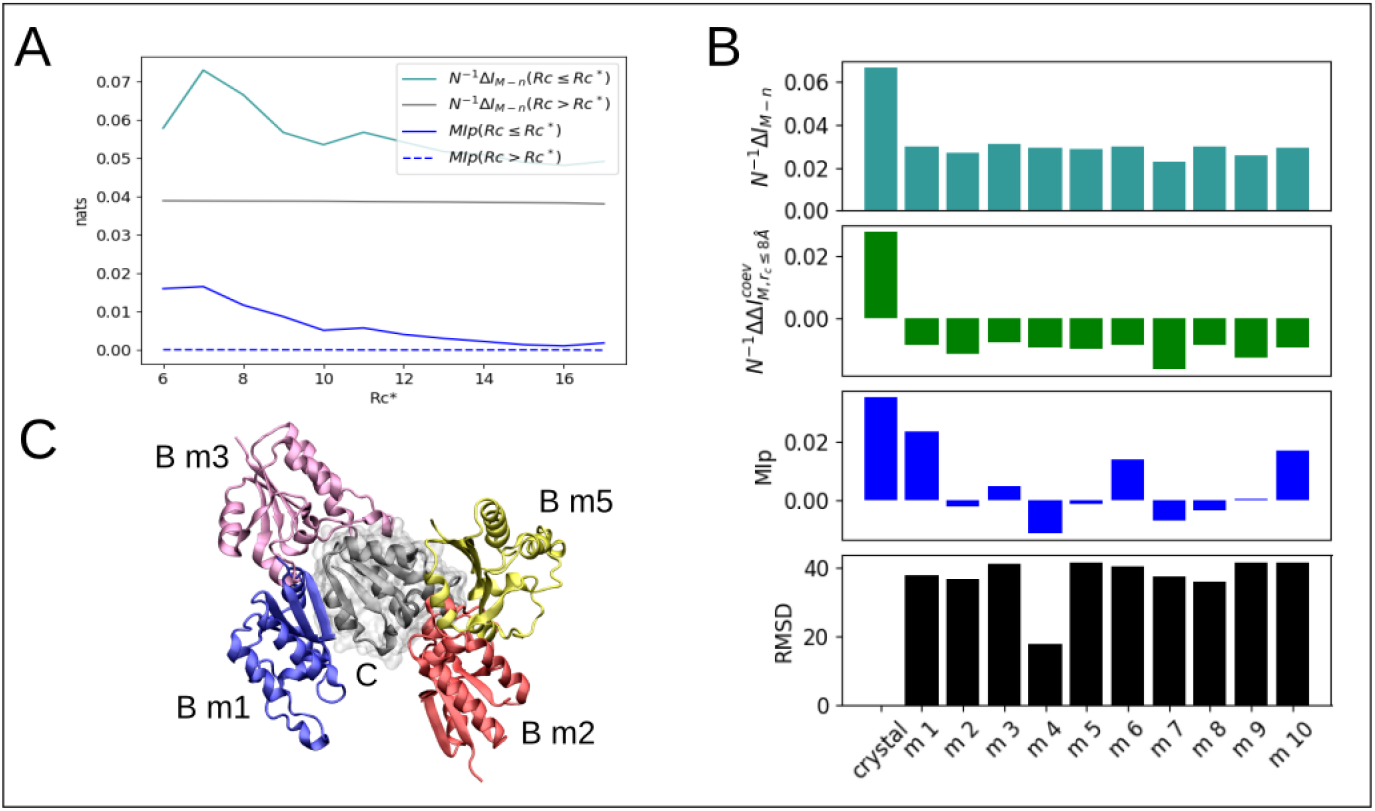
Dependence with contact definition 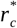 and docking decoys. (A) 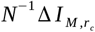 and 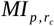, at various 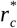. (B) 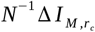 (turquoise), 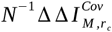 (green), 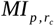 (blue) at alternative interfaces generated by docking – only physically coupled amino acids as defined for *r*_*c*_≤8.0 Å were included in the calculations. Black bars represent the root-mean-square deviation (RMSD) between the native bound structure and docking decoys as generated by GRAMMX (28). (C) Illustration of four docking decoys of chain B in the protein complex TusBCD (chain C is shown in gray).

At 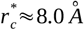, there is still a clear dependence of the per-contact mutual information gap 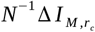, with binding modes or docking decoys of proteins *A* and *B*. Shown in Fig. 3B, the per-contact mutual information gap reaches a maximum at the bound configuration of *A* and *B* (RMSD = 0.0 Å), meaning that 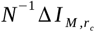, embodies coevolutive pressures in the native amino acids contacts beyond their accessibility at the molecular surface of proteins. The conclusion is further supported by noticing that the isolated coevolutive content 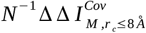 for the bound configuration of *A* and *B* or the associated mutual information *MI*_*p*_ subtracted from structural-functional relationships are larger than the very same estimates for any docking decoys.

## CONCLUDING REMARKS

Overall, molecular coevolution as the maintenance of the binding free-energy of interacting proteins leads their physically coupled amino-acids to store the largest per-contact mutual information 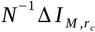, at 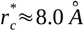, with a significant fraction of the information resulting from coevolutive sources alone. In the present formulation, coupled amino acids are related to the smallest degeneracy ω_*S*_ of native-like MSA models and, therefore, to the largest expectation fraction of PPI matches 〈ϵ〉_*S*_ across such models. These findings hold against any other definition of protein contacts, either across a variety of limitrophe distances *r*_*c*_ discriminating coupled and uncoupled amino acids or alternative binding interfaces in docking decoys. Although presented for the protein complex Tus-BCD, results and discussion also extent to other protein systems as shown in supplementary figures S1 through S4 (*cf.* SI).

Advances in PPI prediction (14–18) are highly welcome in the contexts of paralog matching, host-pathogen PPI network prediction and interacting protein families prediction. Recent studies suggest strategies like maximizing the interfamily coevolutionary signal (14), iterative paralog matching based on sequence “energies” (15) and expectation–maximization (18), which have been capable of accurately matching paralogs for some study cases. Despite these advances, the problem of PPI prediction remains unsolved for sequence ensembles in general, especially for proteins that coevolve in independent genomes though likely resulting from the same free-energy constraints - examples are phage proteins and bacterial receptors, pathogen and host-cell proteins, neurotoxins and ion channels, to mention a few. Far from being trivial, PPI prediction from primary sequences alone is highly complex only solvable within a certain accuracy to date. In this regard, our findings reveal the largest amount of non-stochastic information 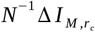 available in primary sequences under coevolution to discriminate MSA models with the largest expectation fraction of PPIs matches 〈ϵ〉_*s*_. Optimality of 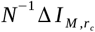 and 〈ϵ〉_*S*_ might be therefore of practical relevance in search of more effective heuristics to resolve PPIs from arbitrary multi-sequence alignments. When compared to evolutive sources, that information is the strongest signal to characterize protein interactions derived from cospeciation and likely, the unique indication in case of coevolution without cospeciation as non-stochastic information of uncoupled amino acids must vanish in independent proteins.

We believe the results are of broad interest as the stability principles of protein systems under coevolution must be universal, either under cospeciation or in independent genomes. We therefore anticipate that decomposition of evolutive and coevolutive information imprinted in physically-coupled and uncoupled amino acids and evaluation of their potential utility in resolving MSA models in terms of degeneracy and fraction of PPI matches should guide new developments in the field, aiming at characterizing protein interactions in general.

## ACKNOWLEDGEMENTS

Comments of Antônio Francisco P. de Araújo. Fernando Melo and Michael Klein on the manuscript are gratefully acknowledged. The research was supported in part by the Brazilian Agencies CNPq, CAPES and FAPDF under Grants 305008/2015-3, 23038.010052/2013-95 and 193.001.202/2016. WT thanks CAPES for doctoral fellowship to MA and CP.

## Supporting Information

### Derivation of Main Text Equations

Consider two proteins *A* and *B* that interact via formation of *i*=1,…, *N* independent amino-acid contacts at the molecular level. Proteins *A* and *B* are assumed to evolve throughout *M* distinct coevolution processes *z* described by the stochastic variable *Z* with probability mass function ρ(*z*), ∀ *z* ∈{1,…, *M*}. Given any specific process *z*, their interacting amino-acid sequences are respectively described by two *N*-length blocks of discrete stochastic variables (*X*_1_, …, *X*_*N*_) and (*Y*_1_, …, *Y*_*N*_) with probability mass functions {ρ(*x*_1_,…, *x*_*N*_),ρ(*y*_1_,…, *y*_*N*_), ρ(*x*_1_,…, *x*_*N*_, *y*_1_,…, *y*_*N*_∣*z*)} such that

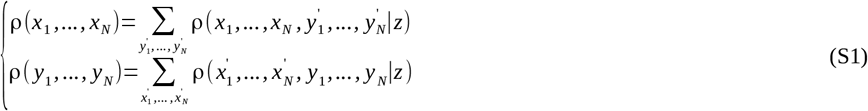

and

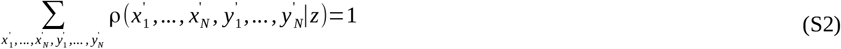

over every joint sequence {*x*_1_, …, *x*_*N*_, *y*_1_, …, *y*_*N*_}_*|χ|*_^*2N*^ defined in the alphabet χ of size |χ|.

Under these considerations, we are interested in quantifying the amount of information that protein *A* stores about the interacting amino-acids of protein *B* conditional to any given coevolution process. As made explicit in eq. [1], we are particularly interested in the situation in which marginals of the *N*-block variables {ρ(*x*_1_,…, *x*_*N*_),ρ(*y*_1_,…, *y*_*N*_)} are independent of process *z* meaning that, for a fixed sequence composition of proteins *A* and *B* only their joint distribution depends on coevolution. Furthermore, by assuming *N*-independent contacts, we want that information to be quantified for the least-constrained model ρ^*^(*x*_1_, …, *x*_*N*_, *y*_1_, …, *y*_*N*_|*Z*) that maximizes the conditional joint entropy between *A* and *B*-that condition ensures the mutual information to be written exactly, in terms of the individual contributions of contacts *i*.

Given its entropy-maximization property (1), ρ^*^(*X*_1_, …, *X*_*N*_, *Y*_1_, …, *Y*_*N*_|*Z*) factorizes into the conditional joint distributions of individual contacts *i*

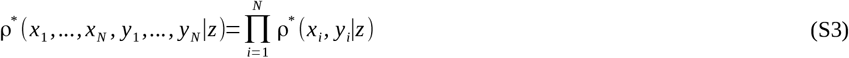

such that

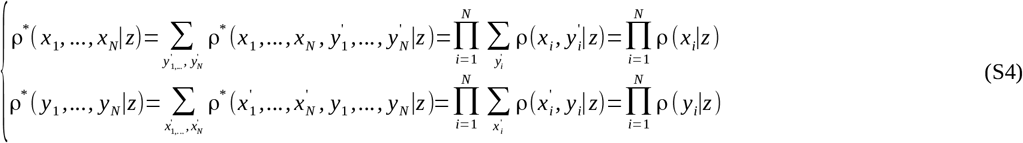

are marginals for any specific *N*-block sequence of proteins *A* and *B*. Eq. [S3] ensures the conditional joint entropy to be written extensively in terms of entropic contributions of contact *i*

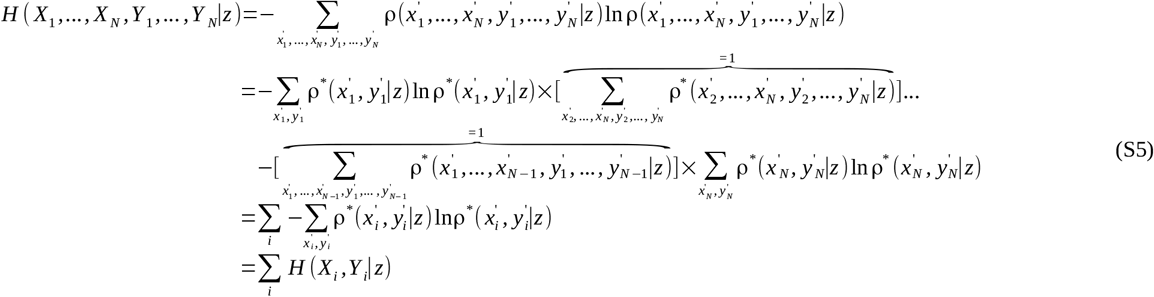

given that

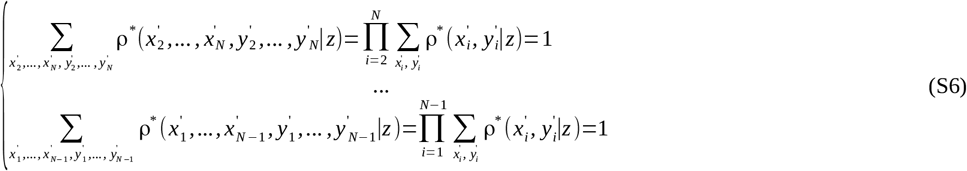

are normalized probabilities of 2(*N* −1)-block sequences. The consequence for the conditional entropy of the individual block variables is then clear

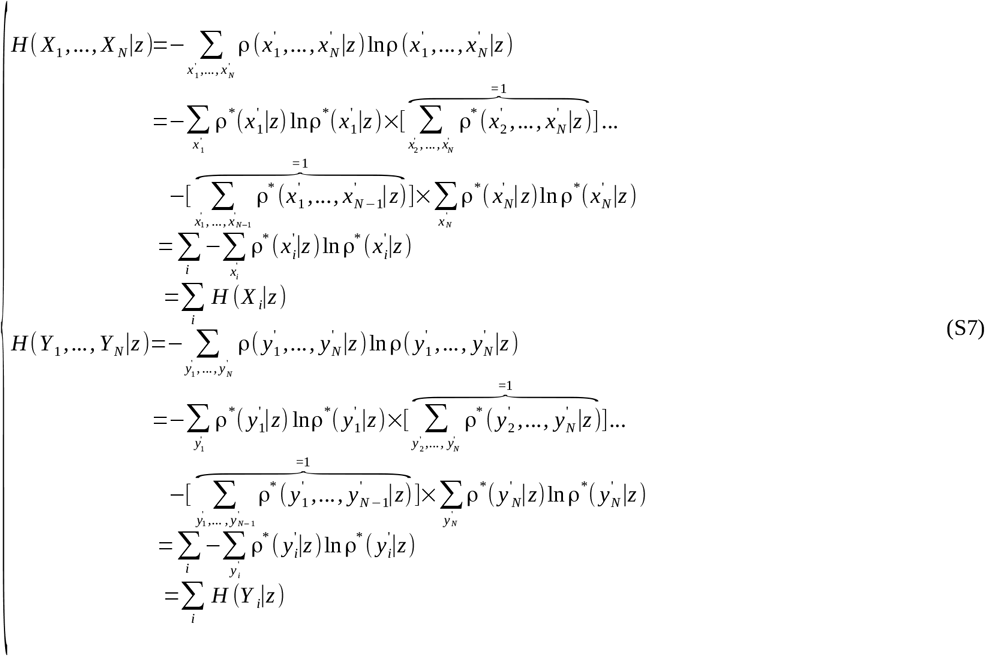

where

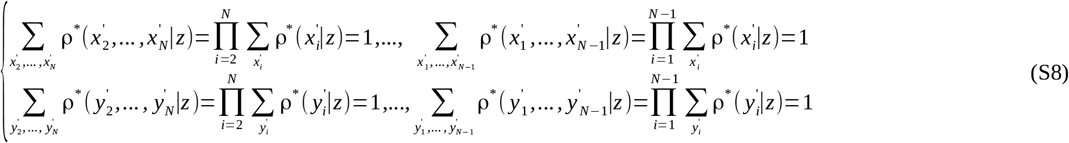

are normalized probabilities of (*N* −1)-block sequences.

Throughout any specific coevolution process *z*, the amount of information that protein *A* stores about the interacting amino-acids of protein *B* is given by the conditional mutual information *I*(*X*_1_,…, *X*_*N*_; *Y*_1_,…, *Y*_*N*_∣*z*) between the stochastic variables (*X*_1_,…, *X*_*N*_) and (*Y*_*1*_,…, *Y*_*N*_).

The expectation value of *I*(*X*^*N*^;*Y*^*N*^∣*z*) across the entire distribution of *M*! distinct coevolution processes reads as

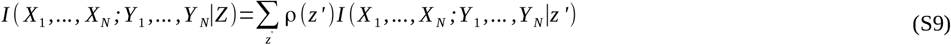

the mutual information between the block variables conditionally to the discrete stochastic variable *Z*. Eq. [S9] can be rewritten

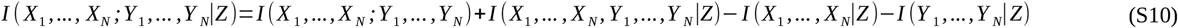

in terms of the information entropies

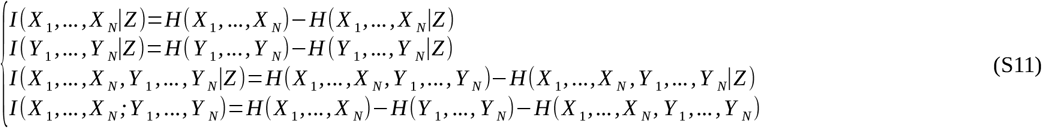

associated with single and joint probability distributions {ρ^*^(*x*_*1*_,…*x*_*N*_|*Z*), ρ^*^(*y*_*1*_,…*y*_*N*_|*Z*), ρ^*^(*x*_*1*_,…*x*_*N*_, *y*_*1*_,…*y*_*N*_|*Z*)} in eq. [S3 and S4]. For the condition in eq. [S1]

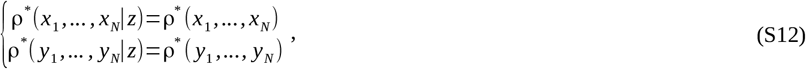

the information entropy of either block variables *H* (*X*_1_,…, *X*_*N*_∣*Z*) and *H* (*Y*_1_,…,*Y*_*N*_∣*Z*) are independent of *Z*

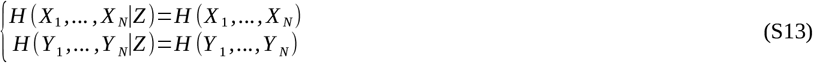

thus simplifying eq. [S10]

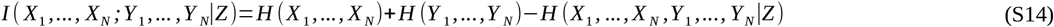

into the joint entropy differences between (*X*_1_,…, *X*_*N*_) and (*Y*_1_,…,*Y*_*N*_) when unconditionally and conditionally dependent on *Z*. From eq. [S5, S7 and S13], the conditional mutual information then rewrites

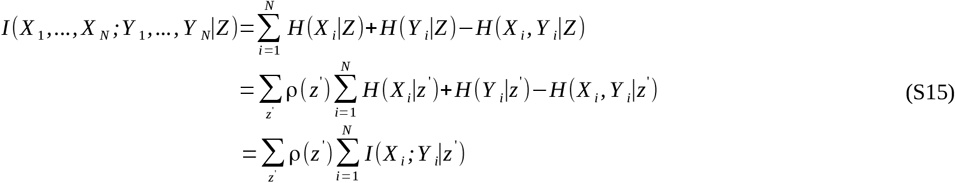

implying

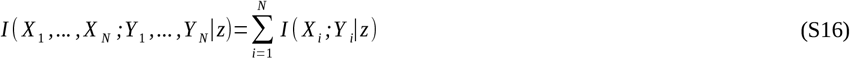

as a direct consequence of eq. [S9].

---

**Fig. S1.**
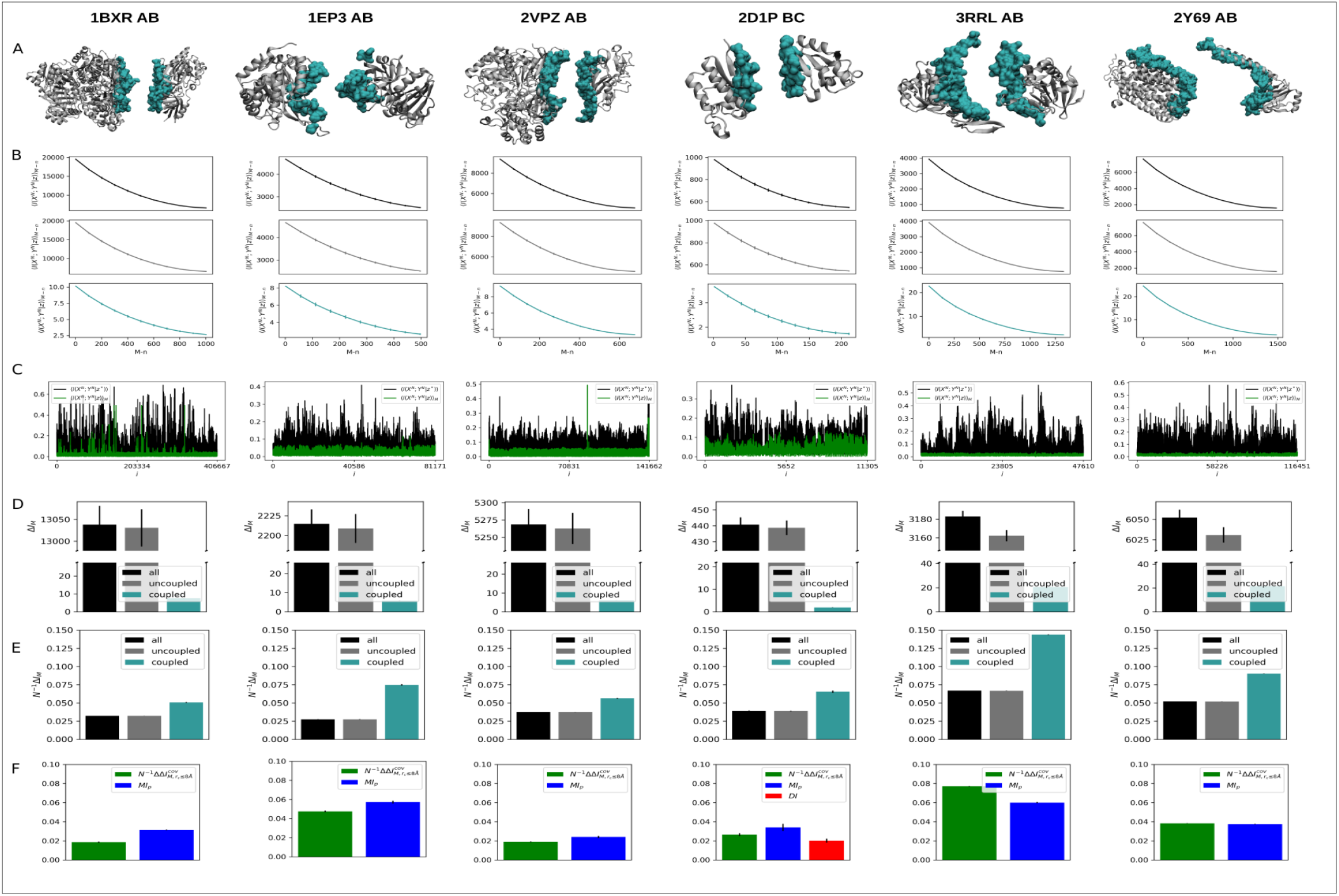
Informational analysis of six protein-protein complexes used in Baker and coworkers.(2) (A) Three-dimensional representation of stochastic variables *X*^*N*^ and *Y*^*N*^ as defined from physically coupled amino acids at short-range cutoff distances *r*_*c*_≤8.0 Å (turquoise) and physically uncoupled amino-acids at long-range cutoff distances *r*_*c*_>8.0 Å (gray). (B) Conditional mutual information 〈 *I*(*X*^*N*^; *Y*^*N*^∣*z*)〉_*M −n*_ as a function of the number *M* − *n* of randomly paired proteins in the reference MSA. 〈*I*(*X*^*N*^; *Y*^*N*^∣*z*)〉_*M −n*_ are expectation values estimated from a generated ensemble of ~ 100 MSA models. (C) Conditional mutual information as a function of protein contact *i*. Mutual information *I*(*X*_*i*_; *Y*_*i*_∣*z*\*) for the reference alignment (black) is systematically larger than 〈 *I*(*X*_*i*_; *Y*_*i*_∣*z*)〉_*M*_ for scrambled models (green) along every contact *i*. (D) Mutual information gap Δ *I*_*M*_ between reference and 100 random models featuring *M* randomly paired sequences. (E) Per-contact mutual information gap 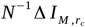. (F) Mutual information decomposition 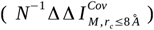 and comparison with functional mutual information 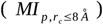 and direct information 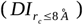. In C, D, E and F error bars correspond to standard deviations.

**Fig. S2.**
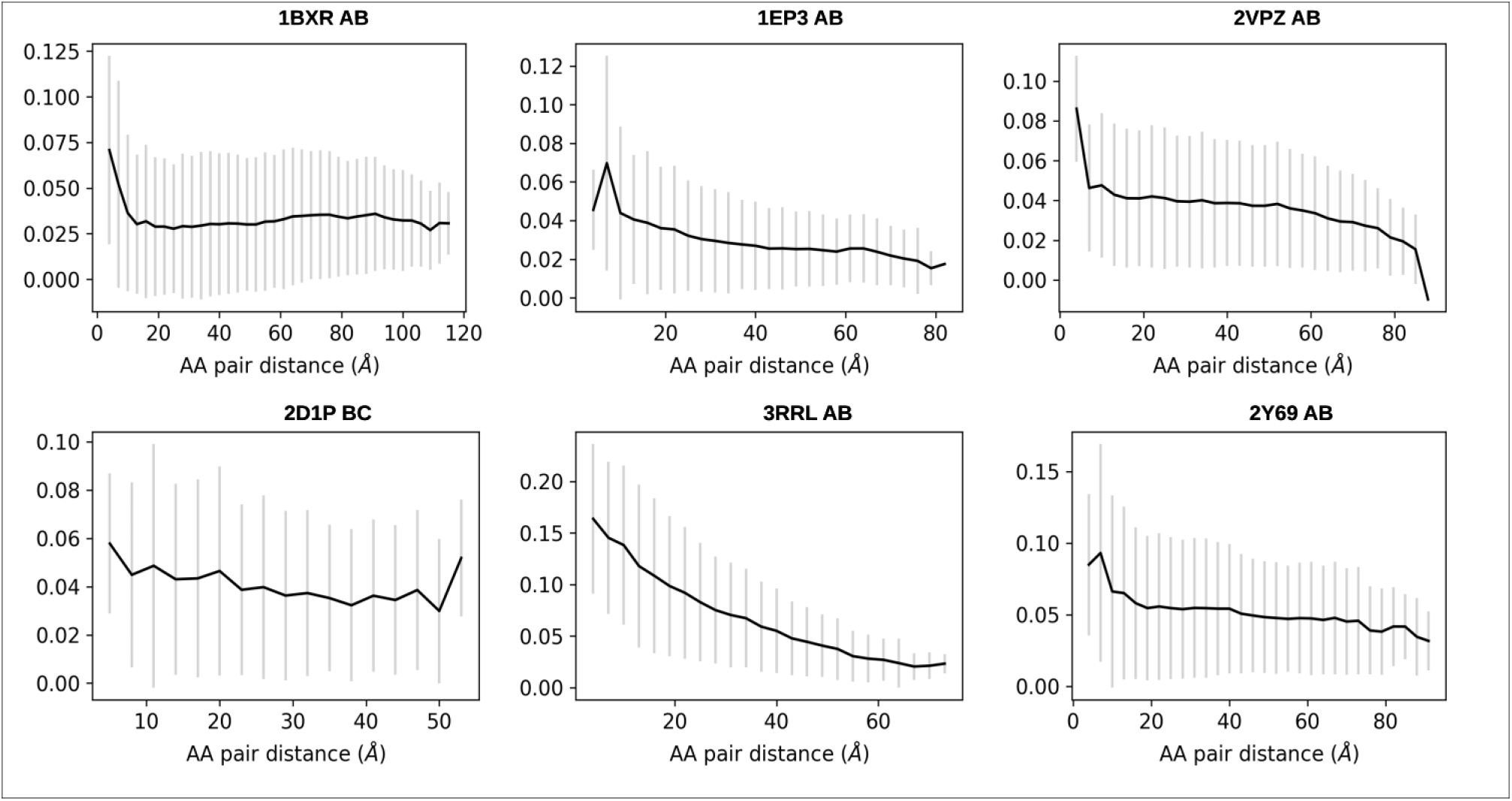
Information gap Δ *I*_*M*_ profile as a function of amino-acid (AA) pair distances. Shown are average values and the associated standard deviations (error bars) of Δ *I*_*M*_ at various pair distances. The profile shows few larger values of Δ *I*_*M*_ at short distances in contrast to many smaller ones at long distances.

**Fig. S3.**
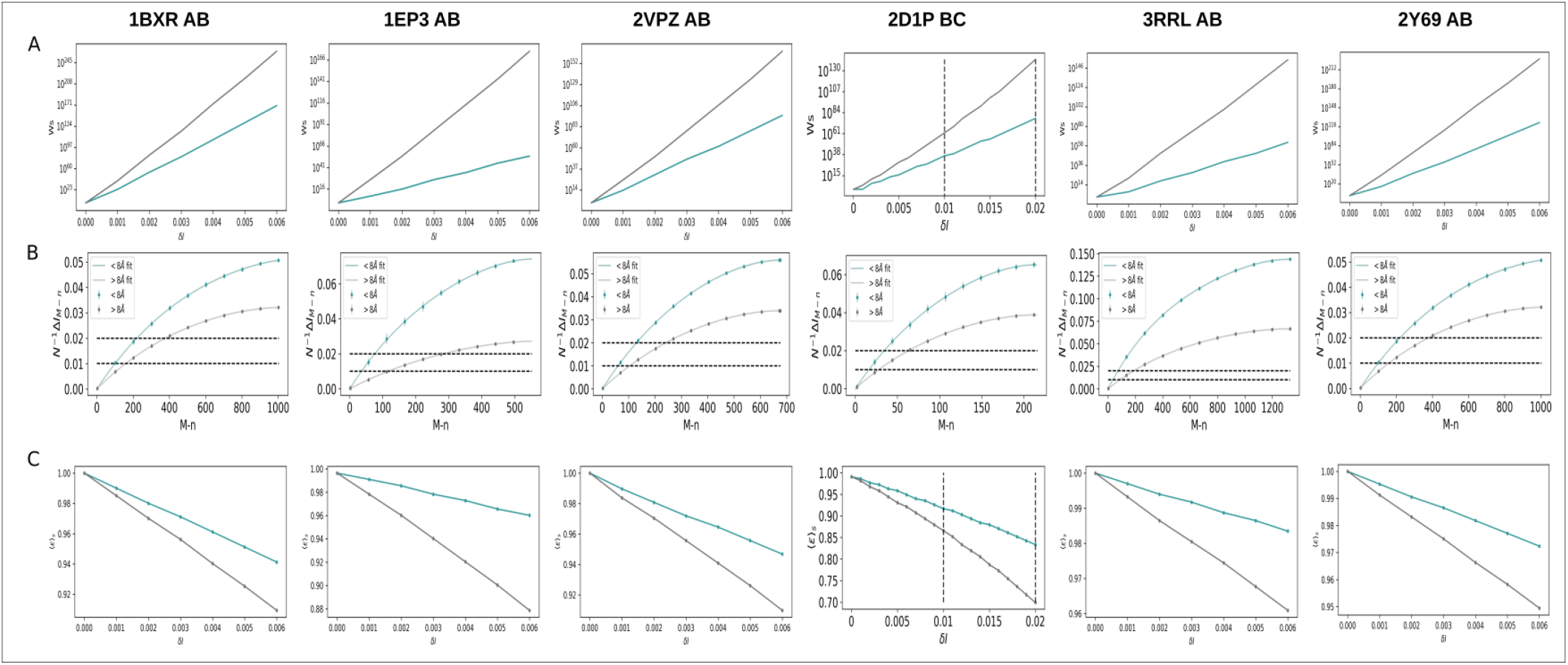
Degeneracy and error analysis for *X*^*N*^ and *Y*^*N*^ involving interacting amino acids at short-range distances *r*_*c*_≤8.0 Å (blue), long-range distances *r*_*c*_>8.0 Å (red), or both (green). (A) Total number ω_*S*_ of native-like models at various resolutions δ*I*. (B) Per-contact gaps of mutual information 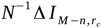 as a function of the number *M* − *n* of randomly paired sequences in the reference alignment. Error bars correspond to standard deviations. (C) Expectation values 〈ϵ〉_*S*_ for the fraction of sequence matches at various resolutions δ*I*.

**Fig. S4.**
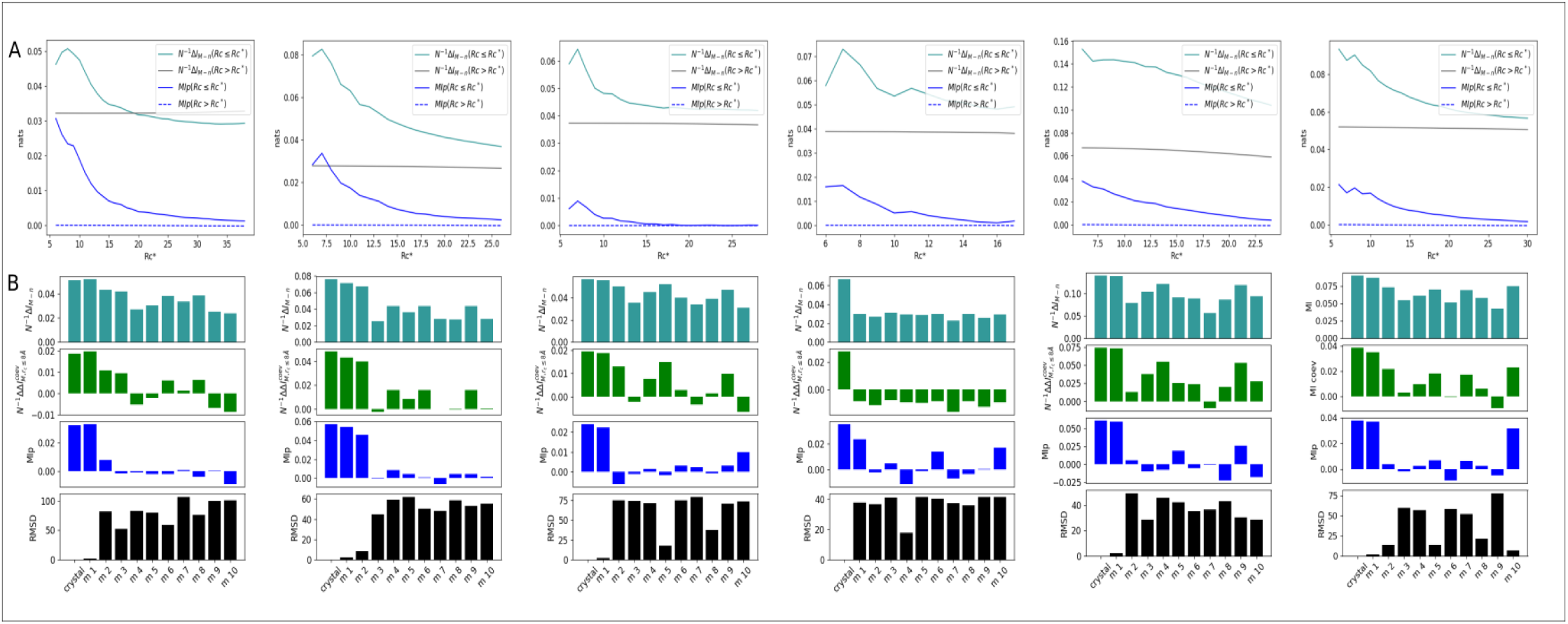
Dependence with contact definition 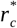 and docking decoys. (A) 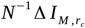 and 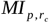 at various 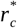. (B) 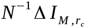 (turquoise), 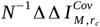(green), 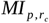 (blue) at alternative interfaces generated by docking – only physically coupled amino acids as defined for *r*_*c*_≤8.0 Å were included in the calculations. Black bars represent the root-mean-square deviation (RMSD) between the native bound structure and docking decoys.

**Table-S1.** Protein system *A* and *B* considered in the study.

| **Complex description** | **PDB ID** | **Protein A** | **Protein B** | **M** | **MSA length** |
| --- | --- | --- | --- | --- | --- |
| Carbamoyl Phosphate Synthetase | 1BXR | **Chain A:** Carbamoyl-Phosphate Synthetase large subunit | **Chain B:** Carbamoyl-Phosphate Synthetase small subunit | 1004 | 1452 |
| Lactococcus Lactis Dihydroorotate Dehydrogenase B. | 1EP3 | **Chain A:** Dihydroorotate Dehydrogenase B (PYRD Subunit) | **Chain B:** Dihydroorotate Dehydrogenase B (pyrk Subunit) | 552 | 572 |
| Polysulfide reductase native structure | 2VPZ | **Chain A:** THIOSULFATE REDUCTASE | **Chain B:** NRFC PROTEIN | 676 | 927 |
| heterohexameric TusBCD proteins | 2D1P | **Chain B:** Hypothetical UPF0116 protein yheM | **Chain C:** Hypothetical protein yheL | 216 | 214 |
| 3-oxoadipate coA-transferase | 3RRL | **Chain A:** Succinyl-CoA:3-ketoacid-coenzyme A transferase subunit A | **Chain B:** Succinyl-CoA:3-ketoacid-coenzyme A transferase subunit B | 1330 | 437 |
| Bovine heart cytochrome c oxidase | 2Y69 | **Chain A:** Cytochrome C Oxidase Subunit 1 | **Chain B:** CYTOCHROME C OXIDASE SUBUNIT 2 | 1484 | 740 |

**Table-S2.**
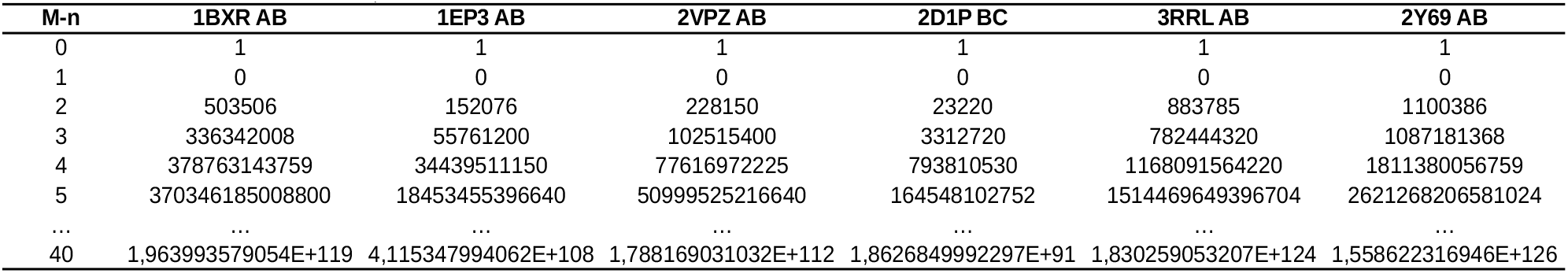
*Rencontres* numbers ω_*M, n*_ as a function of the number *M* − *n* of randomly paired sequences in the reference alignment 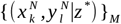.

| Table-S2. *Rencontres* numbers $\omega_{M,n}$ as a function of the number $M-n$ of randomly paired sequences in the reference alignment $\{(x_i^N, y_i^N \vert z^*)\}_M$. | | | | | | |
| --- | --- | --- | --- | --- | --- | --- |
| M-n | 1BXR AB | 1EP3 AB | 2VPZ AB | 2D1P BC | 3RRL AB | 2Y69 AB |
| 0 | 1 | 1 | 1 | 1 | 1 | 1 |
| 1 | 0 | 0 | 0 | 0 | 0 | 0 |
| 2 | 503506 | 152076 | 228150 | 23220 | 883785 | 1100386 |
| 3 | 336342008 | 55761200 | 102515400 | 3312720 | 782444320 | 1087181368 |
| 4 | 378763143759 | 34439511150 | 77616972225 | 793810530 | 1168091564220 | 1811380056759 |
| 5 | 370346185008800 | 18453455396640 | 50999525216640 | 164548102752 | 1514469649396704 | 2621268206581024 |
| ... | ... | ... | ... | ... | ... | ... |
| 40 | 1,963993579054E+119 | 4,115347994062E+108 | 1,788169031032E+112 | 1,8626849992297E+91 | 1,830259053207E+124 | 1,558622316946E+126 |

